# CRISPR loss of function screening to identify genes involved in human primordial germ cell-like cells development

**DOI:** 10.1101/2022.05.23.493064

**Authors:** Young Sun Hwang, M. Andrés Blanco, Kotaro Sasaki

## Abstract

Despite our increasing knowledge of molecular mechanisms guiding various aspects of human reproduction, those underlying human primordial germ cell (PGC) development remain largely unknown. Here, we conducted custom CRISPR screening in an *in vitro* system of hPGC-like cells (hPGCLCs) to identify genes required for acquisition and maintenance of PGC fate in humans. Amongst our candidates, we identified TCL1A, an AKT coactivator. Functional assessment in our *in vitro* hPGCLCs system revealed that TCL1A played a critical role in later stages of hPGCLC development. Moreover, we found that TCL1A loss reduced AKT-mTOR signaling, downregulated expression of genes related to translational control, and subsequently led to a reduction in global protein synthesis and proliferation. Together, our study identifies novel regulators critical for hPGCLC development and demonstrates the importance of translational control in human reproduction.

## INTRODUCTION

Germ cells, including spermatozoa and oocytes, are critical for transmission of genetic and epigenetic information to the next generation. Primordial germ cells (PGCs), which are the early-stage precursors of spermatozoa and oocytes, are specified during early embryonic development through dynamic processes initiated by inductive signals from neighboring tissues. In humans, abnormal germline development leads to a variety of medical conditions, including infertility and congenital diseases. Thus, a comprehensive understanding of the mechanisms regulating germ cell development has critical significance for human health.

Mammalian germ cell development has been extensively studied in mice (Saitou & Yamaji, 2012). However, early embryogenesis differs between mouse and humans (Rossant, 2015), raising concerns as to the translational relevance of such findings to human germ cell development. Indeed, analysis of the specification of PGCs in cynomolgus monkeys (*Macaca fascicularis*), which exhibit a planar structure of peri-implantation embryo, supports the idea that primate PGC specification differs substantially from that in mice with respect to both their origin and developmental trajectories (Sasaki *et al*, 2016). However, analyses of mechanisms underlying germ cell development in humans are hampered by the technical and ethical difficulties in accessing scarce developing germ cells that are only transiently present in human fetal material. Notably, recent advances in an *in vitro* model of human PGCs, termed PGC-like cells (hPGCLCs), derived from human ESCs or iPSCs (Irie *et al*, 2015; Sasaki *et al*, 2015), have provided a foundation for investigating underlying mechanisms that is not restricted by the accessibility and ethical difficulties of research in intact human embryos.

CRISPR loss-of-function genetic screening has been used to discover novel molecular regulators of a wide range of cellular phenotypes in a variety of biological contexts (Shalem *et al*, 2014; Wang *et al*, 2014), including PGCLC specification from mouse ESCs (Hackett *et al*, 2018). However, to date no functional screens for regulators of human germline development and specification have been reported. Here, we carried out a customized CRISPR screen in an *in vitro* system of hPGCLCs derived from a highly quantifiable reporter human inducible pluripotent stem cell (hiPSC) line. In addition, we further elucidated the role of a novel candidate, the AKT coactivator TCL1A, in promoting hPGCLC development.

## RESULTS

### Generation of inducible Cas9 expressing hiPSCs (17C-2 iCas9)

To enable CRISPR loss-of-function genetic screening for regulators of hPGCLC specification and development, we first established an inducible Cas9 (iCas9) hiPSC cell line in which gene editing would occur following hPGCLC induction, but not in the hiPSC state. For this, we utilized a *piggyBac* vector (PB-iCas9-Neo), which encodes both a FLAG-tagged Cas9 and reverse tetracycline-controlled transactivator (rtTA) to confer Doxycycline (Dox)-responsive Cas9 expression, and a Transposase vector for vector integration (**Fig. 1A, MATERIALS AND METHODS**). These vectors were introduced into *TFAP2C-2A-EGFP* and *DDX4/hVH-2A-tdTomato* (AGVT) hiPSCs (585B1 1375, XY), in which EGFP driven by *TFAP2C* promoter (AG) marks hPGCLC and tdTomato driven by *DDX4* promoter (VT) marks gonadal stage germ cells (Hwang *et al*, 2020; Sasaki *et al*., 2015; Yamashiro *et al*, 2018). We isolated seven individual clones and first examined the expression of *rtTA* and *Cas9* by qPCR with or without Doxycycline (Dox) treatment (**Fig. S1A**). As expected, *rtTA* was stably expressed in all clones, regardless of Dox concentration. We selected clone 17C-2, which had the highest Dox-inducible expression of *Cas9,* for further characterization prior to CRISPR screening. We confirmed that clone 17C-2 bore Cas9 expression cassettes and heterozygous AGVT alleles (**Fig. S1B**). Importantly, 17C-2 hiPSCs could be stably maintained in round and tightly packed colonies with a normal karyotype (**Fig. 1B**, **S1C**). We also confirmed that 17C-2 hiPSCs were competent to differentiate into iMeLCs and then AG^+^ hPGCLCs with an induction rate of 38.0% (**Fig. S1D, E**) (Sasaki *et al*., 2015). We next examined the expression of key marker genes during PGCLC induction from 17C-2 hiPSCs (**Fig. S1F**). As expected, pluripotency markers such as *POU5F1* and *NANOG* were consistently highly expressed in the course of induction, whereas *SOX2* expression was limited to hiPSC and iMeLC stages. *EOMES* was specifically expressed in iMeLC stage, and PGC markers such as *BLIMP1*, *SOX17*, *TFAP2C* and *NANOS3* were largely increased in PGCLCs compared to hiPSCs and iMeLCs.

**Fig. 1.**
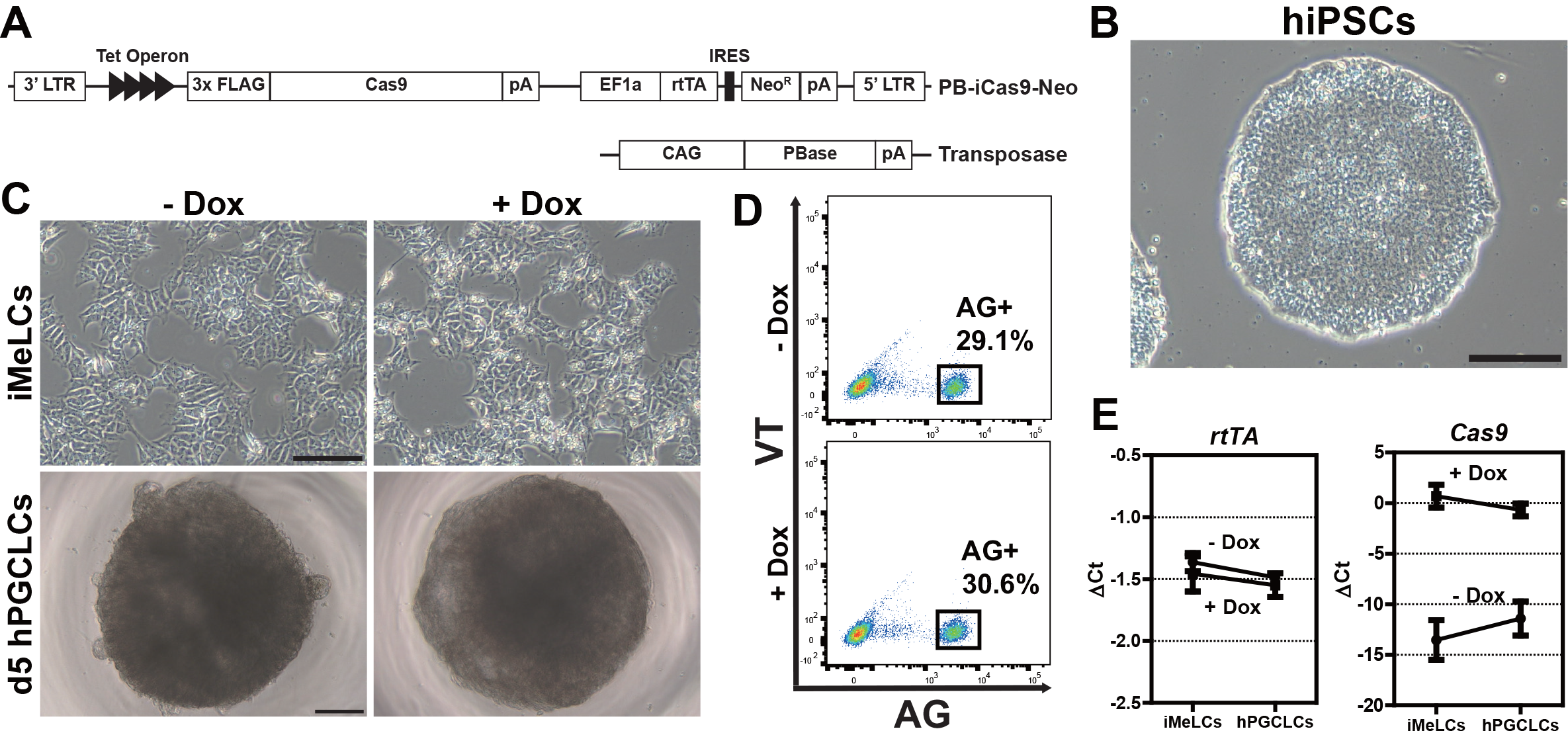
Establishment of inducible Cas9 (iCas9)-expressing hiPSC line. **(A)** Diagram of *piggyBac*-based Cas9 expression under a tet-On inducible system with a PBase (transposase) helper vector. **(B)** A phase-contrast image of 17C-2 (iCas9) hiPSCs. Bar, 200 µm. **(C)** Phase-contrast images of iCas9 hiPSC-derived iMeLCs (top) and day 5 floating aggregates containing hPGCLCs (bottom). – Dox, without Dox; + Dox, treated with 1 µg/ml doxycycline. Bars, 200 µm. **(D)** Fluorescence-activated cell sorting (FACS) analysis of day 5 hPGCLCs derived from iCas9 hiPSCs. AG; TFAP2C-2A-EGFP, VT; DDX4-2A-tdTomato. **(E)** rtTA (top) and Cas9 (bottom) expression during hPGCLC induction, as measured by qPCR. For each gene examined, the ΔCt from the average Ct values of the two independent housekeeping genes *ARBP* and *PPIA* (set as 0) were calculated and plotted. Error bars indicate the standard deviation (SD) of biological replicates.

Prior to undertaking the loss of function CRISPR screen, it was important to determine if Dox treatment affected hPGCLC induction in 17C-2 hiPSCs. Notably, the morphologies of iMeLCs and d5 hPGCLCs (**Fig. 1C**) and the induction efficiency of AG^+^ hPGCLCs (29.1% in -Dox vs. 30.6% in +Dox) were comparable in the presence and absence of Dox (**Fig. 1D**). *rtTA* expression level was also unaffected by Dox during the induction (**Fig. 1E**). Importantly, *Cas9* expression was negligible without Dox, but was highly induced upon Dox treatment in both iMeLCs and d5 hPGCLCs (**Fig. 1E**). Thus, we conclude that the inducible *Cas9*-expressing 17C-2 hiPSCs are highly competent to differentiate into germline, despite high levels of Cas9 expression in response to Dox treatment during hPGCLC induction.

### CRISPR screen during the hPGCLC development *in vitro*

To carry out the loss of function CRISPR screen, we next generated 17C-2 hiPSCs containing a custom CRISPR library. Due to technical hurdles in cell culturing, and the difficulty of generating enough cells for high throughput approaches, we opted to screen a focused, sub-genome scale custom sgRNA library. To best enable identification of hPGCLC regulators, we generated a library focusing on genes that were differentially expressed as iPSC differentiated into oogonia-like cells (Yamashiro *et al*., 2018). These 422 coding-genes were targeted with 5 sgRNAs per gene, and 50 non-targeting control sgRNAs were added for a final library size of 2,208 sgRNAs encoded in the pLentiGuid-Puro vector (**Table S1**).

To optimize the screen, we first determined the lentiviral titer required to achieve a multiplicity of infection (MOI) of 0.3. After 2 days of puromycin selection, we chose 8 μl of lentiviral supernatant per well, which resulted in a 33.3% survival rate (**Fig. S2A**). After **<u>17C-2</u>**-derived **<u>c</u>**ustom **<u>C</u>**RISPR library (17C-2-CC) hiPSCs were generated by lentiviral transduction and selection in large scale, we induced hPGCLCs via iMeLCs, to which Dox was added (**Fig. 2A** and **Fig. S2B**). 17C-2-CC still had a high competency to differentiate into hPGCLCs with a rate of 35.2% of AG^+^ hPGCLCs at day 5 (**Fig. 2B**).

**Fig. 2.**
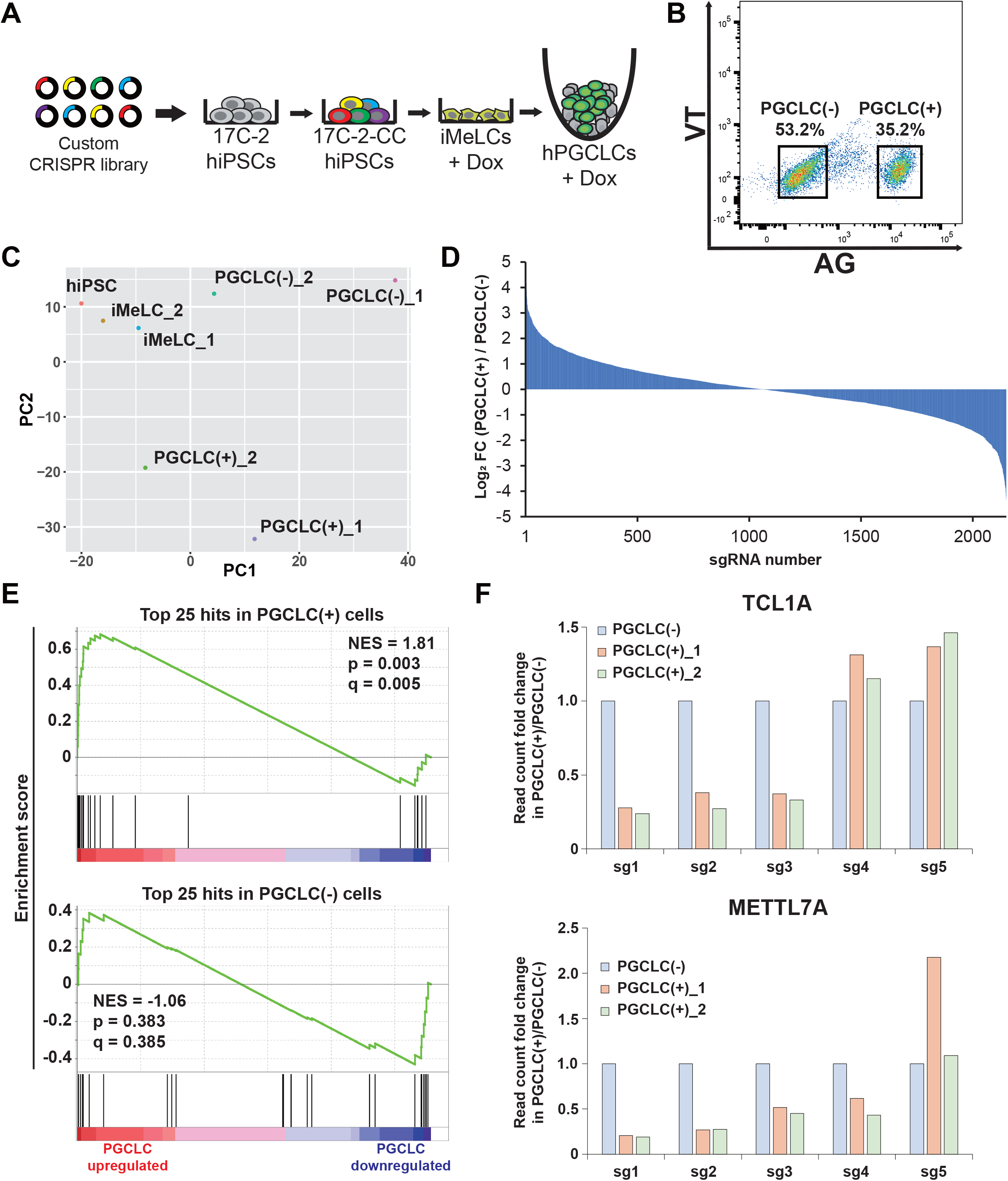
A CRISPR screen for regulators of hPGCLC development using a custom CRISPR library. **(A)** Diagram showing the strategy for CRISPR-based genetic screen during hPGCLC induction *in vitro*. **(B)** FACS sorting of PGCLC(-); AG- and PGCLC(+); AG+ at day 5 of induction. **(C)** PCA of all samples in the screen. **(D)** log_2_ fold change of all sgRNAs in the custom CRISPR library in PGCLC(+) vs. PGCLC(-) cell populations. **(E)** Gene sets consisting of the top 25 hits in PGCLC(+) cells (top) or top 25 hits in PGCLC(-) cells tested for enrichment in RNA-seq of genes most upregulated in PGCLC cells. **(F)** Normalized read counts of sgRNAs targeting *TCL1A* (top) and *METTL7A* (bottom) in PGCLC(+) vs. PGCLC(-) populations in screen replicates. Read counts in in PGCLC(-) cells set to 1.0.

To identify genes potentially driving the hPGCLC state, we isolated Dox-treated AG^+^ hPGCLCs and AG^-^ cells from 17C-2-CC after induction and compared them to untreated hiPSCs and iMeLCs samples. In this screen, sgRNAs depleted from the AG^+^ fraction would represent genes potentially driving the hPGCLC state. Genomic DNA from two biologically replicate screens was then harvested and PCR amplicons of sgRNA barcode abundances were quantified via next generation sequencing. Encouragingly, all samples had similar overall read count distributions (**Fig. S2C**), and replicates had high global similarity (r = 0.92, 0.76, and 0.86 for correlation between replicates of iMeLCs, AG^+^, and AG^-^ samples, respectively) (**Fig. S2D**). PCA revealed a conserved trajectory from hiPSCs to iMeLCs to hPGCLCs between replicates (**Fig. 2C**). Baseline samples (hiPSCs and iMeLCs) had very low gini indices, suggesting initial uniform library representation, and gini indices approximately doubled in AG^+^ and AG^-^ cell fractions (**Table S2**), consistent with selection of variant populations. The distribution of sgRNA log-fold changes was similar in the AG^+^ enriched and AG^-^ enriched sgRNAs (**Fig. 2D**), suggesting a roughly equivalent magnitude of selection in both cell populations. Using the MAGeCK algorithm (Li *et al*, 2014) to call hits on merged replicates, 25 genes scored as significantly depleted from the AG^+^ population at p < 0.05. Among the top hits was *SOX17,* and near-hits included *TFAP2C,* both of which are well-known drivers of the hPGCLC state (**Fig. S2E**). To further confirm the biological validity of the hit list, we assessed the RNA-seq data for enrichment of the top 25 hits, and, as a control, the bottom 25 genes (those with sgRNAs most enriched in AG^+^ cells) in hPGCLCs vs. iMeLCs via GSEA. Consistent with the successful identification of genes whose knockout prevented transition to the hPGCLC state, the top 25 hits were strongly enriched in hPGCLCs compared to iMeLCs, while the bottom 25 genes showed no significant enrichment (**Fig. 2E**). Together, these results suggest that our screen, which to our knowledge is the first of its kind in human germline, was both technically and biologically successful.

### Analysis of hPGCLC induction efficiency by *TCL1A* and *METTL7A* knockouts

To identify candidates for further evaluation, we considered genes that scored at p < 0.05 or were in the top 5 in log fold change in either or both screen replicates (**Table S3**). From this list we selected *TCL1A* and *METTL7A* (**Fig. 2F**) for in-depth validation of functionality in hPGCLC specification, as neither has been linked to germ cell function. TCL1A has an oncogenic role in leukemic T-cell (Russo *et al*, 1989; Virgilio *et al*, 1994) and can act as an AKT coactivator (Laine *et al*, 2000; Pekarsky *et al*, 2000). Its sole MTCP1 domain is encoded by three of its four exons (**Fig. S3A**). METTL7A, a putative m^6^RNA methyltransferase (Wang *et al*, 2022) has two domains – a signal peptide and a methyltransferase domain – that are encoded by its two exons (**Fig. S3B**). To knock out expression of these genes, we decided to remove the ATG start codon and a portion of the domain by two pairs of nCas9 (nickase), respectively (**Fig. S3A, B**). To target each gene, pairs of nickases were cloned into mCherry-expressing vectors and introduced into 1375 (WT) hiPSCs. Two days later, mCherry^high+^ cells (2.26% of *TCL1A* KO cells and 1.86% of *METTL7A* KO cells) were isolated via FACS (**Fig. S3C**). Single clones were expanded (**Fig. S3D**) and screened for the desired deletions via PCR (**Fig. S3E**). Clonal *TCL1A* KO (T1) and *METTL7A* KO (M4) cell lines with confirmed deletions by sequencing (**Fig. S3F**) were then used for subsequent analysis of hPGCLC induction efficiency.

The *TCL1A* and *METT7A* KO lines were induced into hPGCLCs through iMeLCs (**Fig. S3G, H**) and analyzed by flow cytometry. 5 days after induction, we observed a dramatic reduction in induction efficiency in *TCL1A* KO cells, which yielded 8.46% AG^+^ cells (compared to 25.6% WT cells). *METTL7A* KO cells had a more modest, but significant reduction in induction efficiency, generating 17.7% AG^+^ cells (**Figs. S3I and S3J**). Additionally, the number of AG^+^ cells per aggregate was reduced in *TCL1A* (526) and *METTL7A* (1,584) KO cells compared to WT (2,850) (**Fig. S3K**). Thus, these individual knockout approaches validate CRISPR screen for regulators of hPGCLC development. Further, these data suggest that loss-of-function of TCL1A, and to a more moderate extent METTL7A, significantly impairs human germ cell development.

### TCL1A is required for hPGCLC development post-specification

As the phenotype of *TCL1A* KO cells was more severe than that of *METTL7A* KO cells, we further investigated the effect of *TCL1A* KO on hPGCLC development by tracing hPGCLC specification and progression over 6 days (**Fig. 3A, B**). At day 2, the rate and the number of AG^+^ cells per aggregate was similar in KO (17.6%; 1443) and WT (15.5%, 1215) cells (**Fig. 3B-D**). However, at day 4 a significant difference emerged (WT 30.0%, 2176; KO 17.3%, 1191, \*\**P* = 0.0012 (% AG (+) cells)), and at day 6 there was a striking reduction in the number of AG^+^ cells per aggregate in *TCL1A* KO cells compared to WT (WT 20.7%, 2843; KO 16.4%, 545, respectively, \**P* = 0.0230 (AG (+) cells / aggregate)), with *TCL1A* KO cell numbers further decreasing from day 4 to day 6 (**Fig. 3B-D**). This trend suggests that *TCL1A* deletion blocks hPGCLC development after specification and results in their gradual loss over time.

**Fig. 3.**
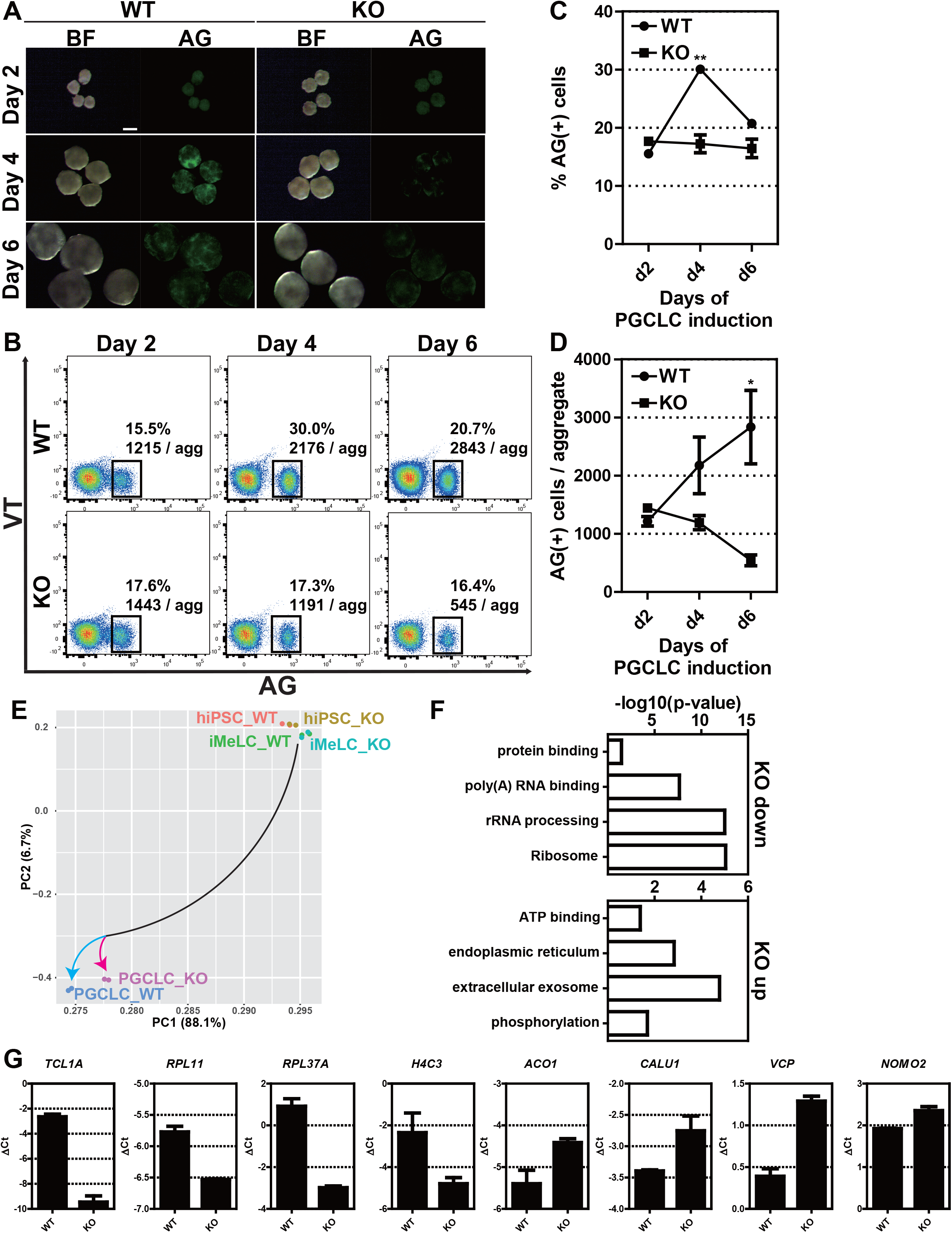
TCL1A is critical for hPGCLC induction. **(A)** Bright field (BF) and fluorescence (AG) images of floating aggregates at days 2, 4 and 6 after hPGCLC induction from wild-type (WT, left) or *TCL1A* knockout (KO, right) hiPSCs. **(B)** FACS analysis of AGVT expression during hPGCLC induction in WT (left) and KO lines (right). Boxes denote AG (+) cells, and their average percentage and the number of AG (+) cells per aggregate from three independent experiments. **(C, D)** The percentage of AG (+) cells (**C**) and the number of AG (+) cells per aggregate (**D**) in WT (circles) or KO hPGCLCs (squares) at the indicated time points. Error bars indicate SD of biological triplicates. Statistically significant differences between WT and KO on each day were identified with a two-tailed *t*-test. \*\**P* = 0.0012 (% AG (+) at d4), \**P* = 0.0230 (AG (+) cells/aggregate at d6) **(E)** PCA of transcriptomes (two independent experiments) of hiPSCs, iMeLCs and day 4 hPGCLCs in WT and KO lines. Color coding indicates the cell types. **(F)** GO analysis of DEGs between WT and KO hPGCLCs. Representative genes in each GO category are shown. **(G)** Gene expression validation of DEGs between WT and KO hPGCLCs by qPCR. For each gene examined, the ΔCt values from the average Ct values of the two independent housekeeping genes *ARBP* and *PPIA* (set as 0) were calculated and plotted. Error bars indicate SD of biological replicates.

To further determine how *TCL1A* deletion blocks hPGCLC development, we performed RNA-seq using hiPSCs, iMeLCs and d4 AG^+^ cells (PGCLCs) and analyzed their global transcriptomes by principal component analysis (PCA) (**Fig. 3E**). Consistent with our functional analysis, transcriptomic features in KO hiPSCs and iMeLCs were highly comparable to those of WT. Developmental trajectories starting from hiPSCs and iMeLCs to hPGCLCs undertook a similar course, but the global transcriptome of PGCLC_KO diverged from that of PGCLC_WT (**Fig. 3E**). Based on the profiles, we identified differentially expressed genes (DEGs) in PGCLC_KO relative to PGCLC_WT (FDR-adjusted *p*-value ≤ 0.05). We found 19 down-regulated and 39 up-regulated DEGs in PGCLC_KO (**Table S4**). Gene ontology (GO) and Kyoto Encyclopedia of Genes and Genomes (KEGG) terms revealed distinct functional classifications for up- and down-regulated genes in PGCLC_KO (**Fig. 3F**, **Table S5** and **Table S6**, *p*-value ≤ 0.05). For the down-regulated genes, enriched gene sets were those involved in “Ribosome (hsa03010)”, “rRNA processing (GO:0006364)”, “poly(A) RNA binding (GO:0044822)” and “protein binding (GO:0005515)” (**Fig. 3F**, **Table S5**). In particular, genes related to Ribosome and rRNA processing, such as seven 60S ribosomal proteins (RPs) (*RPL32*, *RPL34*, *RPL23*, *RPL11*, *RPL37A*, *RPL14* and *RPL19*) and two 40S RPs (*RPS8* and *RPS11*) and a ribosomal protein lateral stalk subunit P2 (*RPLP2*), were downregulated in *TCL1A* KO hPGCLCs (Table S5). By contrast, functional GO terms in up-regulated genes included “extracellular exosome (GO:0070062)”, “endoplasmic reticulum (GO:0005783)”, “ATP binding (GO:0005524)” and “phosphorylation (GO:0016310)” (**Fig. 3F**, **Table S6**). Specifically, *ANKRD13C*, *VCP*, *SEC23IP*, *CALU*, *ACO1*, *DHCR24*, *PLOD1*, *P4HB*, *NOMO2* and *ZMPSTE24* associated with endoplasmic reticulum or endoplasmic reticulum membrane were upregulated in *TCL1A* KO hPGCLCs. We also confirmed gene expression changes including down-regulated genes such as *TCL1A*, *RPL11*, *RPL37A* and *H4C3*, and up regulation of *ACO1*, *CALU1*, *VCP* and *NOMO2* by qPCR (**Fig. 3G**). Based on the significant downregulation of ribosome-associated genes, these data support a potential role for *TCL1A-*mediated translational control of hPGCLC development.

### TCL1A regulation of hPGCLC development through AKT signaling

TCL1A is a coactivator of AKT signaling (Laine *et al*., 2000; Pekarsky *et al*., 2000), highlighting a potential role for AKT in hPGCLC development. To determine if AKT signaling was affected by *TCL1A* knockout in hPGCLCs, we quantified Pan-AKT and phosphorylated AKT (p-AKT) by immunocytochemistry (**Fig. 4A**), as there was insufficient protein from hPGCLCs for Western blot analysis. All hPGCLCs in both WT and KO were positive for AKT protein and its expression level was unchanged after *TCL1A* knockout (**Fig. 4A, C**). However, the intensity of p-AKT was lower in KO (*p* = 0.0001) and the number of p-AKT cells were also reduced in KO (56.5%, 35/62; *p* = 0.0018) compared to WT (27.5%, 17/62) (**Fig. 4C, D**). These results indicate that TCL1A acts as an AKT coactivator in hPGCLCs.

**Fig. 4.**
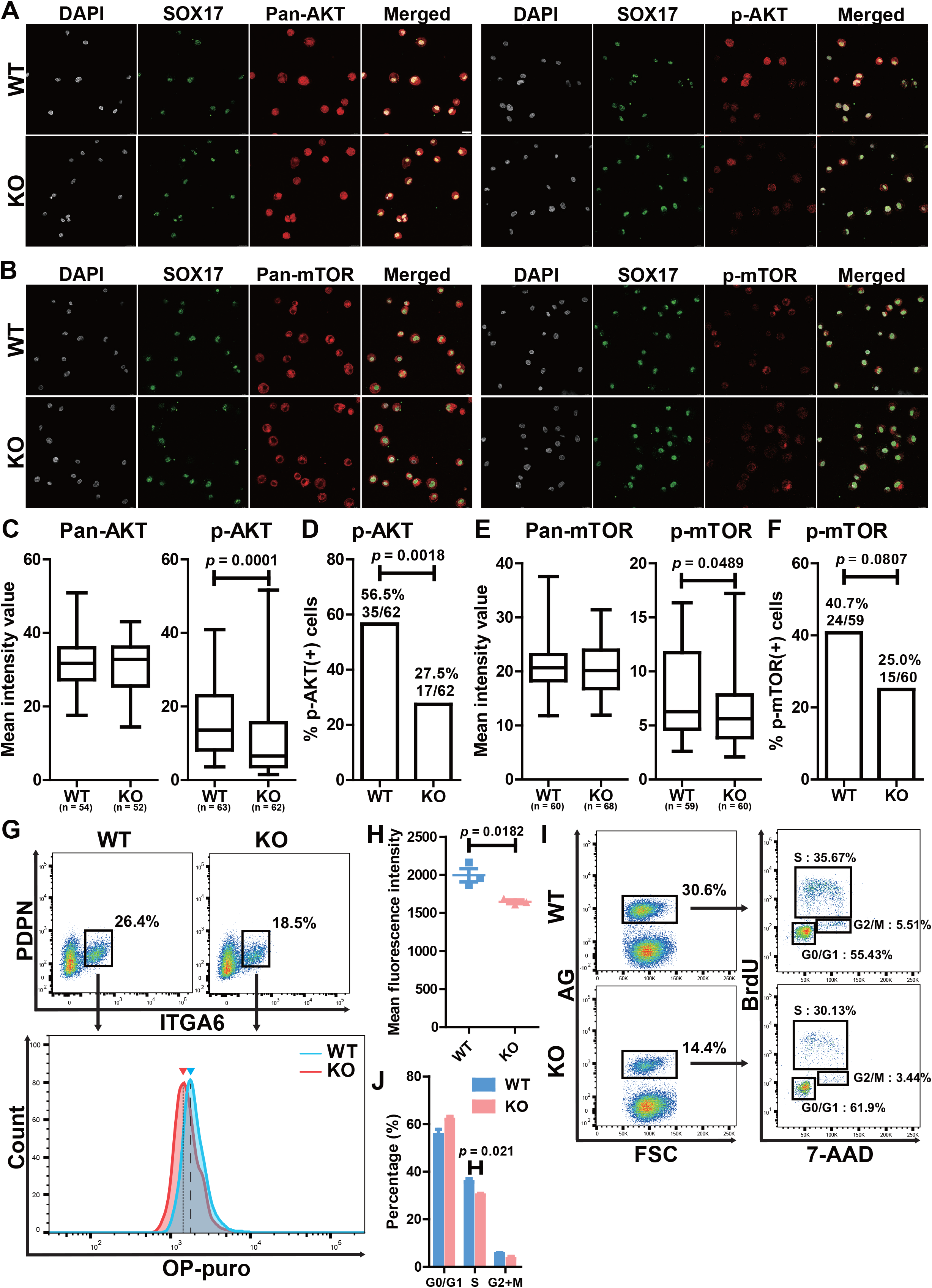
Functional analysis of TCL1A in hPGCLCs. (A, B) Immunocytochemistry analysis of 1375 (WT) and *TCL1A* knockout (KO) hPGCLCs for DAPI (white), SOX17 (green), pan-AKT, phospho (p)-AKT, pan-mTOR or p-mTOR (red). Bar, 20 µm. **(C, E)** Box plot quantification of the mean intensity values of the indicated markers, Center line, median; box limits, upper and lower quartiles; whiskers, 1.5× interquartile range. Statistically significant differences between WT and KO were identified with Mann– Whitney *U* test. *P* = 0.0001 (p-AKT), 0.0489 (p-mTOR). n, number of SOX17^+^ cells counted. **(D, F)** Bar graphs showing the proportions of the indicated phosphorylated proteins in SOX17^+^ cells. Statistically significant differences were identified with Fisher’s exact test. *P* = 0.0018 (p-AKT), 0.0807 (p-mTOR) **(G)** Protein synthesis assay in WT and KO hPGCLCs, on the basis of OP-puro incorporation. (top) FACS analysis of ITGA6 and PDPN expression. Boxed areas indicate the percentage of ITGA6^+^PDPN^+^ hPGCLCs. (bottom) Representative FACS histograms of OP-puro incorporation in ITGA6^+^PDPN^+^ hPGCLCs derived from WT or KO lines. The blue arrowhead and dashed line indicate the mean fluorescence intensity (MFI) of WT hPGCLCs, and the red arrowhead and dotted line indicate the MFI of KO hPGCLCs. **(H)** Scatter dot plot of MFI of OP-puro, measured by FACS. Each dot (blue, WT; red, KO) represents three independent experiments, and lines indicate mean MFI values. Statistically significant differences were identified with two-tailed *t*-test. *P* = 0.0182. Error bars indicate SD of biological triplicates. **(I)** Cell cycle analysis of WT and KO hPGCLCs by BrdU and 7-AAD corporation. (left) FACS analysis of AG expression. The percentages of AG (+) cells (highlighted in boxes) are shown. (right) BrdU and 7-AAD analysis of AG (+) cells. The average percentage of cells in the indicated cell cycle stages are shown. **(J)** Bar graph of the percentage of each cell cycle stage. Statistically significant differences were identified with two-tailed *t*-test. *P* = 0.021 (S phase). Error bars indicate SD of biological triplicates.

Our findings that the percentage and the number of hPGCLCs was decreased by *TCL1A* knockout was associated with downregulation of Ribosome and rRNA processing by RNA-seq, supporting a role for TCL1A in translational regulation of hPGCLC development (**Fig. 3A-D, F**). As the mTOR pathway downstream of AKT signaling regulates cell growth, proliferation, and survival through translational control (Ruggero & Sonenberg, 2005), we next assessed mTOR signaling by immunocytochemistry (**Fig. 4B**). While mTOR protein was globally expressed in hPGCLCs and its expression level was comparable between WT and KO (**Fig. 4E**), the expression level of p-mTOR was down-regulated by *TCL1A* knockout (**Fig. 4E**; *p* = 0.0489) and the number of p-mTOR positive cells was reduced in KO hPGCLCs compared to WT, although not significantly (**Fig. 4F**; *p* = 0.0807). Therefore, mTOR signaling downstream of AKT is altered by *TCL1A* knockout in hPGCLCs.

To determine if defective mTOR signaling following *TCL1A* knockout impacts protein synthesis in hPGCLCs, we next utilized a flow cytometric O-propargyl-puromycin (OP-puro) incorporation assay. After selection of hPGCLCs by surface markers that included PDPN and ITGA6, we found that the peak and level of OP-puro intensity in KO was significantly decreased compared to WT (**Fig. 4G, H**; *p* = 0.0182). Flow cytometric cell cycle analysis of hPGCLCs by BrdU and 7-AAD incorporation revealed that the proportion of G0/G1 (55.43% to 61.9%) phase was increased, whereas that of S (35.67% to 30.13%) and G2/M (5.51% to 3.44%) phase was decreased in KO relative to WT (**Fig. 4I, J**), although only S phase was significantly different (*p* = 0.021). Thus, these results demonstrate that global protein synthesis level and cell cycle were altered through AKT-mTOR signaling pathways in hPGCLC after *TCL1A* knockout.

## DISCUSSION

Using a novel Dox-inducible Cas9-expressing hiPSC line carrying fluorescent reporters specific for human germline development (iCas9_AGVT_17C-2) (**Fig. 1**), we provide the quantitative and qualitative functional analysis of the effect of gene knockout at specific stages of human germ cell development. Notably, this strategy could easily be combined with xrOvary or xrTestis to examine more differentiated cell types (Hwang *et al*., 2020; Yamashiro *et al*., 2018). Using this novel platform, we conducted the first targeted CRISPR screening designed to identify novel factors important for hPGCLC development (**Fig. 2**). Our identification of numerous candidates that could potentially contribute to human germ cell development, one of which is described further below, provide essential insight into hPGCLC development and identify multiple pathways for future mechanistic studies.

Although key transcription factors for human germline specification, such as SOX17 (Irie *et al*., 2015; Kojima *et al*, 2017), BLIMP1 (Sasaki *et al*., 2015), and TFAP2C (Kojima *et al*., 2017; Pastor *et al*, 2018) have been functionally assessed, the mechanistic basis for hPGCLC-specification remains largely enigmatic. Among candidates identified by CRISPR screen analysis in the current work, we showed that TCL1A plays a role in hPGCLCs development post-specification (**Fig. 3**). Although induction of hPGCLCs progressed normally in *TCL1A* KOs, the number of AG^+^ cells decreased over time (**Fig. 3A-D**). Thus, TCL1A appears important for the proliferation and/or survival of cells already committed to the germline, thus differing from well-known key elements contributing to the early specification of hPGCLCs (Kojima *et al*., 2017; Kojima *et al*, 2021; Sasaki *et al*., 2015).

TCL1A is a proto-oncogene first identified in leukemic T-cells (Russo *et al*., 1989; Virgilio *et al*., 1994) that acts as a coactivator of AKT signaling (Laine *et al*., 2000; Pekarsky *et al*., 2000). In primordial germ cells, AKT appears to be activated by kit ligand (also known as stem cell factor, SCF)-dependent PI3K signaling (De Miguel *et al*, 2002; Farini *et al*, 2007). As shown in **Fig. 4A, C, D**, phosphorylation of AKT was decreased in *TCL1A* KO hPGCLCs. Notably, transcriptome analysis of WT and KO hPGCLCs revealed that TCL1A-AKT signaling likely impacted translation, as GO and KEGG analysis revealed that terms related to translation such as rRNA processing and Ribosome were down-regulated in KO cells while terms related to endoplasmic reticulum were up-regulated (**Fig. 3F**). Interestingly, previous studies revealed that TCL1A was significantly decreased during hPGCLC expansion culture, suggesting that additional metabolic changes contribute to human germ cell proliferation (Murase *et al*, 2020).

mTOR signaling downstream of AKT mediates translational control (Ruggero & Sonenberg, 2005), driving global expression of most RP genes (Cardenas *et al*, 1999; Preiss *et al*, 2003) and modulating the activity of transcription factors, such as TIF-1A, important for Pol I activity and rRNA synthesis (Mayer *et al*, 2004). Consequently, the AKT-mTOR signaling pathway drives ribosome biogenesis and global protein synthesis critical for cell growth, proliferation, and survival. Consistent with these effects, we found that TCL1A-mediated AKT signaling is compromised in hPGCLC KOs and results in downregulation of mTOR signaling (**Fig. 4B, E, F**). Moreover, in hPGCLCs, inhibition of mTOR signaling by mutant TCL1A reduced global level of protein synthesis, likely contributing to decreased cell proliferation and/or survival (**Fig. 4G-J**). Thus, TCL1A-dependent regulation of AKT-mTOR signaling appears to play a critical role in translational control in hPGCLCs, ultimately leading to their decrease in number over time following knockout. Translation of a subset of mRNAs responsible for hPGCLCs development appears to be controlled by mTOR signaling in other cell types as well (Hsieh *et al*, 2012; Thoreen *et al*, 2012), further highlighting the importance of this pathway.

In conclusion, we have generated essential genetic resources critical for further mechanistic study of hPGCLC development and developed a CRISPR screening strategy that will aid in the further identification of genes involved in human germline development. Using this platform, our current study demonstrated the potential importance of TCL1A regulation of AKT-mTOR dependent translational control of cell proliferation and/or survival in hPGCLCs. As translational control of germline stem cells of *Drosophila* (Sanchez *et al*, 2016) has previously been documented, our studies clearly support further studies on the translational regulation of human germ cell development and reproduction.

## MATERIALS AND METHODS

### Culture of hiPSCs

All experiments using human primordial germ cell-like cells (hPGCLCs) induced from human induced pluripotent stem cells (hiPSCs) were approved by the University of Pennsylvania Institutional Review Board (IRB; protocol #832470). hiPSCs were cultured on plates coated with recombinant laminin-511 E8 (iMatrix-511 Silk, Nacalai USA) and maintained under feeder-free conditions in StemFit® Basic04 medium (Ajinomoto) containing 20 ng/ml basic FGF (Peprotech) at 37°C under a 5% CO_2_ atmosphere. Before passaging or the induction of differentiation, hiPSC cultures were treated with a 1:1 mixture of TrypLE Select (Life Technologies) and 0.5 mM EDTA/PBS for 15 min at 37°C to dissociate them into single cells. Subsequently, 10 μM ROCK inhibitor (Y-27632; Tocris) was added in culture medium for 1 day after passaging hiPSCs.

### Generation of inducible Cas9 (iCas9) expressing hiPSC lines

For the construction of an inducible Cas9 expression piggyBac vector (PB-iCas9-Neo), a Gateway entry vector containing Cas9, pENTR-hSpCas9, was cloned into a PB-TA-ERN plasmid (all-in-one piggyBac transposon destination vector for doxycycline (Dox)-inducible expression with constitutive rtTA and neomycin resistance) (Kagawa *et al*, 2019; Kim *et al*, 2016) with Gateway LR Clonase II enzyme mix (Invitrogen). The reactions were performed according to the manufacturer’s protocol. These plasmids were provided by Dr. Knut Woltjen at Kyoto University. Detailed cloning histories are available upon request as Snapgene files. The PB-iCas9-Neo vector (2 µg) and pCAG-PBase vector (1 µg) (transposase expression vector, a gift from Dr. Knut Woltjen) (Kim *et al*., 2016) were introduced into 1 million AGVT hiPSCs (585B1 1375, gift from Dr. Mitinori Saitou, Kyoto University) by electroporation with a NEPA21 Type II electroporator (Nepagene). Single colonies were isolated after selection with neomycin and chosen for expansion and characterization.

For the measurement of rtTA and inducible Cas9 expression, each clone was treated with Dox (Takara) at 0, 250 or 1000 ng/ml for 2 days. Total RNA was extracted with an RNeasy Micro Kit (QIAGEN) according to the manufacturer’s instructions. Total RNA (1 μg) was used as a template for cDNA synthesis with the SuperScript III First-Strand Synthesis System (Invitrogen). The cDNA was serially diluted five-fold, and equal amounts were PCR amplified. Q-PCR was performed with Power SYBR Green PCR Master mix (Life Technologies) and a StepOnePlus™ Real-Time PCR System (Applied Biosystems). The gene expression levels were examined through calculation of ΔCt (on log2 scale) normalized to the average ΔCt values of PPIA and ARBP, with the primer pairs listed in Table S7. The success of the integration of the Cas9 cassette was assessed by PCR of the extracted genomic DNA from each clone. Clone 17C-2 (designated as iCas9_AGVT_17C-2) was selected for iCas9 hiPSCs.

### Karyotyping and G-band analyses

hiPSCs were incubated in culture medium containing 100 ng/ml of KaryoMAX Colcemid solution (Gibco) for 8 h. After dissociation by TrypLE Select, cells were treated with pre-warmed buffered hypotonic solution and incubated for 30 min at 37°C. Cells were then fixed with Carnoy’s solution (3:1 mixture of methanol and acetic acid) and added dropwise onto glass slides for preparation of chromosomal spreads. Karyotypes were first screened by counting the numbers of chromosomes identified by DAPI staining. Cell lines bearing 46 chromosomes were further analyzed on the basis of G-banding by Cell Line Genetics (Madison, WI).

### Induction of hPGCLCs

hPGCLCs were induced from hiPSCs via iMeLCs as described previously (Sasaki *et al*., 2015). For the induction of iMeLCs, hiPSCs were plated at a density of 4×10^4^ to 5×10^4^ cells/cm^2^ onto a human fibronectin (Millipore)-coated 12-well plate in GK15 medium (Life Technologies) with 15% KSR, 0.1 mM NEAA, 2 mM L-glutamine, 1 mM sodium pyruvate and 0.1 mM 2-mercaptoethanol) containing 50 ng/ml of ACTA (R&D Systems), 3 μM CHIR99021 (Tocris Bioscience) and 10 μM of a ROCK inhibitor. After 31 h, iMeLCs were harvested and dissociated into single cells with TrypLE Select. For induction of hPGCLCs, iMeLCs were then plated at 3,500 cells per well into a low-cell-binding V-bottom 96-well plate (Thermo Fisher Scientific) containing GK15 medium supplemented with 200 ng/ml BMP4 (R&D Systems, 314-BP-010), 100 ng/ml SCF (R&D Systems, 255-SC-010), 50 ng/ml EGF (R&D Systems, 236-EG), 10 ng/ml LIF (Millipore, #LIF1010) and 10 μM of Y-27632 (GK15+BSELY). The plates were incubated at 37°C under a 5% CO_2_ atmosphere until use in downstream assays.

### Q-PCR analysis of marker expression

Total RNA from iMeLCs and hPGCLCs was extracted with an RNeasy Micro Kit (QIAGEN) according to the manufacturer’s instructions. The concentration of total RNA was measured with a Qubit 3 fluorometer (Invitrogen). Synthesis and amplification of cDNAs with 1 ng of purified total RNA was performed as previously described (Nakamura *et al*, 2015). Q-PCR on amplified cDNA was performed with Power SYBR Green PCR Master mix (Life Technologies) and a StepOnePlus™ Real-Time PCR System (Applied Biosystems). The gene expression levels were examined by calculation of *ΔC_t_* (on log_2_ scale) normalized to the average *ΔC_t_* values of *PPIA* and *ARBP*, with the primer pairs listed in **Table S7**.

### Fluorescence-activated cell sorting (FACS)

We isolated AG^+^ hPGCLCs from floating aggregates by using FACS. Floating aggregates containing hPGCLCs, which were induced from hiPSCs, were dissociated into single cells through 0.1% trypsin/EDTA treatment for 15 min at 37°C with periodic pipetting. After the reaction was quenched by addition of an equal volume of FBS, cells were resuspended in FACS buffer (0.1% BSA in PBS) and strained through a 70 µm nylon cell strainer (Thermo Fisher Scientific) to remove cell clumps. AG^+^ cells were sorted with a FACSAria Fusion flow cytometer (BD Biosciences) and collected in an microcentrifuge tube containing CELLOTION (Amsbio). All FACS data were collected in FACSDiva v8.0.2 software (BD Biosciences) and analyzed with FlowJo v10.8.1 (BD Biosciences). For flow cytometry analysis, hPGCLCs were dissociated and resuspended in FACS buffer. AG^+^ cells were analyzed with FACSCanto or FACSFortessa (BD Biosciences).

### CRISPR/Cas9 Screen Analysis

A total of 422 genes were selected from 453 previously reported genes variably expressed during the transition of hPGCLC-derived cells in xenogeneic reconstituted ovaries, after removal of pseudogenes and long-non-coding RNAs (Yamashiro *et al*., 2018). A custom CRISPR gRNA library pool containing 2159 gRNAs was designed and synthesized by GenScript (Piscataway, NJ). The PCR products were amplified and ligated into the pLentiGuid-Puro vector through *BsmBI* restriction via GenBuilderᵀᴹ Plus assembly. The library was verified through next generation sequencing by GenScript. For lentivirus packaging, the target vector and pCMV-VSV-G and pCMV-dR8.2 dvpr lentiviral packaging plasmids (Addgene plasmid #8454 and #8455, gifts from Dr. Robert Weinberg, Massachusetts Institute for Technology) were co-transfected into 293T cells (Clontech, 632180) with PEI reagent (Polysciences, 23966-1). Lentiviral particles were collected 48 h and 72 h after transfection and filtered.

For CRISPR screening, 1 million 17C-2 hiPSCs were transduced with 0, 8 or 16 μl of lentivirus supernatant for 6 h in a 6 cm dish. One day after transduction, cells were treated with puromycin for 48 h. Three days after transduction, hiPSCs transduced with each titer were harvested and counted, and an MOI of 0.3 was measured (**Fig. S2A**). The transduction with 8 µl of lentivirus supernatant was identified as the optimal titer to achieve an MOI of 0.3. Three million 17C-2 hiPSCs in three 6 cm dishes were then transduced with 8 µl lentivirus supernatant per dish, selected with puromycin and expanded in bulk; the gRNA fold coverage was maintained, and these cells were designated as 17C-2-CC hiPSCs. iMeLCs and hPGCLCs derived from 17C-2-CC were induced as described above, except that 1 µg/ml Dox was added for induction of Cas9 expression. In every step, the gRNA fold coverage was maintained above 200×. At day 5, floating hPGCLC aggregates, and AG^+^ [designated as PGCLC(+)] and AG^-^ cells [designated as PGCLC(-)] were sorted with a FACSAria Fusion flow cytometer and collected in 15 ml tubes containing CELLOTION. Genomic DNA from hiPSCs, iMeLCs, PGCLC (+) and PGCLC (-) was isolated with a GenElute Mammalian Genomic DNA Miniprep kit (Sigma).

Genomic DNA libraries for sequencing were then prepared according to previously described protocols (Joung *et al*, 2017) maintaining >200X fold coverage. Briefly, PCR-amplified library samples were purified with the QIAquick PCR Purification Kit (Qiagen) followed by gel extraction with the QIAquick Gel Extraction Kit (Qiagen). The barcoded libraries were then pooled at equimolar ratios and sequenced on a NextSeq500/550 instrument (Illumina, 150 cycles High Output kit v2.0) to generate 150-bp single-end reads. MAGeCK software was used for screening analysis (Li *et al*., 2014) using default settings to call hits in PGCLC(+) vs. PGCLC(-) cells, with screen replicates analyzed separately. All computer codes used are available upon request.

### Generation of *TCL1A* and *METTL7A* knockout hiPSC lines

Two pairs of sgRNA sequences before and after the targeted exons of *TCL1A* and *METTL7A* were designed with the Molecular Biology CRISPR design tool (Benchling). Two pairs of sgRNAs targeting *TCL1A* (CGAGTGCCCGACACTCGGGG and GCAAGAGCCAGAGCCTCTCA for pre-exon1, and AGGTACAGCCAGCTTTGGAG and GGCTGTACCTCGATGGTTAA for post-exon1) or *METTL7A* (TACCATCTTTATCCTGAGAC and GTAAGCTCCATTGCTCAGAA for pre-exon1, and CGGGGAGAACATGCACCAGG and AGCGCTCAAACTGCAGGTGT for post-exon1) were selected and cloned into the pX335-U6-Chimeric BB-CBh-hSpCas9n (D10A) SpCas9n-expression vector to generate the sgRNAs/Cas9n vector (Addgene, #42335; (Cong *et al*, 2013) . Four sgRNAs/nCas9 vectors (2.5 µg each) were introduced into 1 million AGVT hiPSCs (1375) by electroporation with a NEPA21 Type II electroporator. At 2 days after transfection, mCherry-expressing cells were sorted into a 96-well plate (single cell/well) with a FACSAria Fusion flow cytometer, and subsequently expanded and genotyped by PCR (**Fig. S3E**). Large deletions in T1 and M4 were further validated by Sanger sequencing (**Fig. S3F**).

### RNA-seq Analysis

Total RNA from hiPSCs, iMeLCs and hPGCLCs (AG^+^) was extracted with an RNeasy Micro Kit, and RNA quality was evaluated with High Sensitivity RNA Screen Tape on an Agilent 4200 TapeStation. The cDNA library was prepared with a Takara SMART-Seq HT kit, and adapters were ligated with a Nextera XT DNA Library Preparation Kit and Index Kit v2 according to the manufacturers’ protocols. AMPure XP beads were used for primer cleanup. Subsequently, 75-base pair reads were sequenced on a NextSeq 500 instrument (Illumina, San Diego, CA)) according to the manufacturer’s protocol.

The quality of FASTQ files was verified with FastQC v0.11.2. All samples were aligned to the human genome, build hg38, with Rsubead aligner v1.24.1(Liao *et al*, 2019). The number of fragments overlapping each Entrez gene were summarized by using featureCounts (Liao *et al*, 2014) and Rsubread’s inbuilt hg38 annotation. Differentially expressed genes (DEGs) were identified with the DESeq2 package with FDR ≤ 0.05 as the significance threshold. DEGs were mapped to GO terms by using DAVID (v6.8) with the background list set to “Homo sapiens” (Huang *et al*, 2007). Only enriched GO terms with *p* < 0.05 were shown.

### Immunofluorescence (IF) analysis

For IF staining of hPGCLCs, day 4 aggregates were dissociated into single cells and spread onto poly-L-lysine-treated Superfrost™ Plus Microscope Slides (Fisher Scientific). The slides were fixed in 4% paraformaldehyde in PBS for 15 min at room temperature (RT), washed three times with PBS for 5 min, permeabilized with 0.2% Triton-X in PBS for 10 min at RT and washed three times with PBS for 5 min. Next, the slides were incubated with primary antibodies to pan-AKT (CST, 4691T), phospho-AKT (Ser473; CST, 4060T), pan-mTOR (CST, 2983T) and phospho-mTOR (Ser2448; CST, 5536T) in blocking solution (1% bovine serum albumin in PBS) overnight at 4℃. After being washed three times with blocking solution for 5 min, the slides were incubated with secondary antibodies in blocking solution and 1 µg/ml of DAPI for 1 hr in the dark at RT. The slides were washed with blocking solution three times for 5 min, then mounted in Vectashield mounting medium (Vector Laboratories) for confocal laser scanning microscopy analysis (Leica, SP5-FLIM inverted). Confocal images were processed in Leica LasX (v3.7.2). The mean fluorescence intensity of each marker in SOX17-positive cells was measured in ImageJ software (National Institutes of Health, Bethesda, MD).

### Cell cycle analysis by BrdU and 7-AAD incorporation

BrdU (1 mM) in GK15 was diluted in GK15+BSELY medium to obtain a final concentration of 20 µM. Then half the medium of hPGCLC cultures in V-bottom plates was changed (final BrdU concentration of 10 µM). BrdU-treated aggregates were incubated at 37°C under 5% CO_2_ in air for 1 h. hPGCLCs were dissociated into single cells, and cell cycle analysis was performed by staining cells with BrdU-APC and 7-AAD with BD Pharmingen™ APC BrdU Flow Kits (552598, BD Biosciences), according to the manufacturer’s instructions. Stained cells were analyzed in FACSFortessa.

### Protein synthesis assay by OP-puro incorporation

O-propargyl-puromycin (OP-puro) labeling was performed with a Protein Synthesis Assay Kit (ab235634; Abcam) according to the manufacturer’s instructions. Protein Label stock (200×) was diluted in GK15+BSELY medium to produce 1× Protein Label solution. Then day 4 aggregates containing hPGCLCs were transferred onto Nunc™ 4-Well Dishes (Thermo Fisher Scientific) containing 1× Protein Label solution. Dishes were incubated at 37°C under 5% CO_2_ in air for 1 h. To identify hPGCLCs, we used surface markers, because GFP signals would have been quenched by the sodium azide treatment during the procedure. hPGCLCs were dissociated into single cells and then stained with Alexa Fluor 647-conjugated rat anti-human PDPN (337007, BioLegend) and BV421-conjugated anti-CD49f (INTEGRINα6) (313623, BioLegend) in FACS buffer for 30 min at RT before fixation. Protein synthesis was detected with FACSFortessa.

## Data Availability

The datasets generated in this study are available at NCBI GEO under the following accession numbers: GSE202996.

## ACKNOWLEDGMENTS

We thank L. King for carefully reviewing the manuscript and providing insightful comments. We thank members of Sasaki and Blanco laboratory for discussion of this study. This work was supported in part by the Open Philanthropy funds from the Silicon Valley Community Foundation (2019-197906) and Good Ventures Foundation (10080664) to K.S.

## AUTHOR CONTRIBUTIONS

K.S. and M.A.B. conceived and supervised the project. Y.S.H. designed and conducted the overall experiments. K.S., M.A.B and Y.S.H wrote the manuscript. M.A.B. contributed to the analysis of CRISPR screening data.

## Figure Legends

**Fig. S1. Generation and selection of iCas9 hiPSC clones, associated with Fig. 1. (A)** Gene expression of *rtTA* (left) and *Cas9* (right) in the indicated hiPSC clones. White, gray and black bars indicate cells treated with Dox at 0, 250 or 1000 ng/ml, respectively. The quantification of the gene expression levels was as shown in Fig. 1E. Error bars indicate SD of technical duplicates. Clone 17C-2 (highlighted in red) was selected for downstream CRISPR screening assays because it showed the highest *Cas9* expression after Dox treatment. **(B)** PCR genotyping of the *TFAP2C-2A-EGFP* (AG) (left), *DDX4-2A-tdTomato* (VT) (middle) and *Cas9* coding sequence (right). Rec., recombined with fluorescent protein; non, non-targeted; yellow arrow, 3 kb; yellow asterisk, 1 kb. **(C)** Representative results of 17C-2 hiPSC karyotype analysis, showing a normal karyotype (46, XY). **(D)** Phase-contrast image of 17C-2 hiPSC-derived iMeLCs. Bar, 200 µm. **(E)** Bright-field image of a day 5 floating aggregate derived from 17C-2 hiPSCs (top) and its FACS plot. Bar, 200 µm. The percentages of AG (+) cells (highlighted in box) are shown. **(F)** Gene expression dynamics of key markers during hPGCLC induction from iCas9 hiPSCs. hPGCLCs were harvested on day 5. Error bars indicate the standard deviation (SD) of technical duplicates. n.d., not detected.

**Fig. S2. Generation of 17C-2 iCas9 hiPSCs containing a custom CRISPR library (17C-2-CC) by lentiviral transduction, associated with Fig. 2. (A)** (left) Phase-contrast images of lenti-viral transduced 17C-2 hiPSCs before (day 1) and after (day 3) puromycin selection. Volumes of virus-containing supernatant added per well (0 µl, 8 µl and 16 µl; virus titer) are indicated. (+), with puromycin; (-), without puromycin. (right) The number of cells counted at day 3 in each titer. 0 (-) is set as 100%. Titer 8 was selected as the optimized titer for a multiplicity of infection (MOI) of 0.3. **(B)** Phase-contrast images of 17C-2-CC hiPSCs (left), iMeLCs (middle) or day 5 floating aggregates containing hPGCLCs derived from 17C-2-CC hiPSCs (right). Bars, 200 µm. Distribution of abundances of normalized sgRNA read counts in all screen samples. Comparison of read counts of all sgRNAs in biological replicates of PGCLC(+) screen samples. **(E)** Normalized read counts of sgRNAs targeting *TFAP2C* (left) and *SOX17* (right) in PGCLC(+) vs. PGCLC(-) populations in screen replicates. Read counts in in PGCLC(-) cells set to 1.0.

**Fig. S3. Knockout scheme of TCL1A and METTL7A mutant hiPSCs and analysis of their induction efficiency of hPGCLCs, associated with Fig. 3. (A, B)** Knockout schemes for human *TCL1A* (A) or *METTL7A* (B) loci. I in A, TCL1_MTCP1 domain; I and II in B, METTL7A signal peptide and Methyltransf_11 domain, respectively. Black boxes indicate exons. Red arrowheads indicate target sites recognized by the pair of Cas9 nickases flanking the start codons and the domain-encoding regions. Arrows indicate primer sites for genotyping. **(C)** FACS sorting of 1375 hiPSCs (WT) on the basis of mCherry expression by px335-derived nickase vectors targeting *TCL1A* (left) and *METTL7A* (right). The percentage of mCherry^high^ cells (highlighted in boxes) and the sorting gates are shown. **(D)** Phase-contrast images of single clonal expansion of *TCL1A-* and *METTL7A*-targeted hiPSCs. Bar, 200 µm. **(E)** PCR genotyping of the large deletion in *TCL1A* (left) and *METTL7A* (right) exons by double pairs of nickase. Yellow asterisk, 500 bp. T1 and M4 (red) clones were selected for further analysis of each knockout line. **(F)** Sanger sequencing results of large deletions in T1 (left) and M4 (right). The thin black line represents a gap between target sites recognized by the pair of Cas9 nickases. **(G, H)** Phase-contrast images of iMeLCs (**G**) and day 5 floating aggregates containing hPGCLCs (**H**) derived from T1 (left) and M4 (right). Bars, 200 µm. **(I)** FACS analysis of day 5 hPGCLCs derived from WT (left), T1 (middle) and M4 (right). Boxes indicate AG (+) cells. The average percentage and number of AG (+) cells per aggregate from two independent experiments are also shown. **(J, K)** Percentage of AG (+) cells (**J**) and number of AG (+) cells per aggregate (**K**) in WT (white), T1 (gray) and M4 (black) in day 5 hPGCLCs. Error bars indicate SD of biological replicates.

